# Evolutionary Analysis of Transcriptional Regulation Mediated by Cdx2 in Rodents

**DOI:** 10.1101/2021.03.01.433326

**Authors:** Weizheng Liang, Guipeng Li, Huanhuan Cui, Yukai Wang, Wencheng Wei, Siyue Sun, Diwen Gan, Rui Chen, Hongyang Yi, Bernhard Schaefke, Yuhui Hu, Qi Zhou, Wei Li, Wei Chen

## Abstract

Differences in gene expression, which can arise from divergence in *cis*-regulatory elements or alterations in transcription factors binding specificity, are one of the most important causes of phenotypic diversity during evolution. By protein sequence analysis, we observed high sequence conservation in the DNA binding domain (DBD) of the transcription factor Cdx2 across many vertebrates, whereas three amino acid changes were exclusively found in mouse Cdx2 (mCdx2), suggesting potential positive selection in the mouse lineage. Multi-omics analyses were then carried out to investigate the effects of these changes. Surprisingly, there were no significant functional differences between mCdx2 and its rat homologue (rCdx2), and none of the three amino acid changes had any impact on its function. Finally, we used rat-mouse allodiploid embryonic stem cells (RMES) to study the *cis* effects of Cdx2-mediated gene regulation between the two rodents. Interestingly, whereas Cdx2 binding is largely divergent between mouse and rat, the transcriptional effect induced by Cdx2 is conserved to a much larger extent.

**Author summary:** Our study 1) represented a first systematic analysis of species-specific adaptation in DNA binding pattern of transcription factor. Although the mouse-specific amino acid changes did not manifest functional impact in our system, several explanations may account for it (See Discussion part for the detail); 2) represented a first study of cis-regulation between two reproductively isolated species by using a novel allodiploid system; 3) demonstrated a higher conservation of transcriptional output than that of DNA binding, suggesting the evolvability/plasticity of the latter; 4) finally provided a rich data resource for Cdx2 mediated regulation, including gene expression, chromatin accessibility and DNA binding etc.

## Introduction

Gene expression refers to the spatiotemporal conversion of information from DNA to functional gene products such as proteins. Knowing how gene expression is regulated is critical for the understanding of development as well as evolution [1]. Indeed, differences in gene expression are considered to be among the most important causes of phenotypic diversity across species [2]. Multiple layers are involved in the regulation of gene expression, of which transcriptional regulation is considered to be a crucial contributor to phenotypic alterations during evolution.

Transcriptional regulation is mediated by the interaction between *cis*-regulatory elements (e.g., promoters and enhancers) and *trans*-factors (e.g., transcription factors (TFs)) [3, 4]. On the one hand, changes in the *cis*-elements located in the vicinity of target genes affect TF binding and/or local chromatin environment, thereby modulating gene expression in one-to-one manner [5]. On the other hand, alterations in *trans*-factors influence the expression of their target genes in a more pleiotropic fashion [3, 6, 7]. Investigation of the *cis-trans* regulatory crosstalk is of great importance for the mechanistic understanding of the genetic basis leading to coordinated phenotypic diversity, and possibly also with disease relevance [4, 6]. Two main strategies have been widely applied to investigate the *cis* and *trans* acting regulatory components in transcriptional regulation: quantitative trait loci (QTL) mapping analysis and F1 hybrid studies. By correlating a measured molecular trait (e.g., gene expression level or TF binding intensity) with genetic variations, expression quantitative trait loci (eQTL) and chromatin immunoprecipitation quantitative trait loci (ChIP-QTL) have been performed to study *cis* and *trans* regulatory divergence in populations [8, 9]. In comparison, the F1 hybrid approach has been used to study *cis* and *trans* regulatory divergence contributing to differences in gene expression between strains of the same species or closely related species. With two alleles sharing the same *trans* environment, allelic differences in the F1 hybrid can be directly interpreted as *cis*-regulatory divergence [10, 11]. By comparing these allele-specific variations with the differences between parental strains or species, the *trans*-component of gene expression differences can be estimated [3, 5, 12, 13]. Whereas gene expression differences between strains of the same species in yeast or Drosophila can be mainly attributed to *trans*-regulatory divergence [3, 14–24], with the proportion of *cis*-divergence increasing with phylogenetic distance, an F1 hybrid study of different *Mus musculus* subspecies exhibited pervasive *cis*-regulatory differences [25]. In line with this finding, divergence of TF binding occupancy could predominately be attributed to *cis*-acting variation in the same F1 hybrid cross [13].

Alteration in *trans*-regulation could arise from either the change in the abundance of the *trans*-regulators or the mutation of their amino acid sequences. In general, evolution of amino acid sequences is much slower than that of non-coding regulatory elements, particularly for the TF DNA binding domains (DBD) [26]. It has been shown that most of the TF orthologs display highly conserved binding specificities between fly and human [27]. This observation is likely due to the fact that changes in TF binding affinities would induce large pleiotropic effects in gene regulation [28]. Therefore, under strong selection constraints, most of the affinity-changing mutations in TF-DBDs would be removed by purifying selection. Alternatively, it is also possible that changes in TF-DBD might have the potential to drive large phenotypic changes if the resulting effects have a net positive effect on the organism’s fitness [29]. If so, species-specific changes in TF-DBD might be positively selected. So far, however, this possibility has been largely unexplored.

In this study, to find a candidate TF which might have undergone adaptive evolution in either the mouse or the rat lineage, we compared the amino acid sequences of TF-DBDs among mouse, rat and human, and searched for the TFs with highly conserved DBD between human and one of the rodent species, but showing multiple amino acid changes in the other rodent species. It turned out that the DBD in caudal-type homeobox 2 (Cdx2) contains three amino acid changes exclusively in mouse. The finding was further substantiated by including 56 species as well as 37 mouse strains in the sequence comparison. Given the established function of Cdx2 in lineage specification and trophectoderm differentiation [30–33], we investigated the potential effect of the three mouse-specific amino acid changes in the mouse embryonic stem cell (mESC) systems. Unexpectedly, we did not observe any significant effects at either DNA binding specificity or target gene expression induced by the three changes. Then, to study the *cis*-regulatory changes in Cdx2-mediated transcriptional regulation between rat and mouse, we analyzed the allele-specific binding of Cdx2 as well as allele-specific transcriptional output induced by Cdx2 in rat-mouse allodiploid embryonic stem cells (RMES) [34]. Interestingly, whereas the Cdx2 binding is largely divergent between mouse and rat, the transcriptional effect induced by Cdx2 is conserved to a much larger extent.

## Results

### Exclusive amino acid changes in the DNA binding domain of mCdx2

The house mouse (*Mus musculus*) and the common rat (*Rattus norvegicus*) are two of the most widely used model organisms in biomedical research. As TF-DNA binding is crucial in gene regulation, we were interested in identifying TFs which might differ in this important aspect between mouse and rat, which might have undergone adaptive evolution in either lineage but are conserved with human in the other. To find such a candidate TF we selected all 1194 TFs with one-to-one orthologs among mouse, rat and human. Systematic analysis of the DNA binding domain (DBD) sequences of these TFs between human, mouse and rat showed the DBD sequence of Cdx2 was identical between human and rat, but possessed three amino acid changes in mouse. Moreover, by analyzing the protein sequences of Cdx2 among four additional model species, surprisingly, we found that these three specific amino acid changes are exclusively present in mouse (Fig 1A). Further comparison of Cdx2 DBD sequences in 56 vertebrate species showed that the DBD is highly conserved in general (S1A Fig) whereas the three amino acid changes were present in all mouse strains with available genome sequences (S1B Fig). Based on this observation, we inferred that the three amino acid changes have occurred on the *Mus* lineage shortly after its divergence from the *Rattus* lineage at about 12.5 to 5 MY (million years) ago (Fig 1B) [35]. Taken together, our evolutionary analyses indicated that mouse Cdx2 had three specific amino acid changes in the DBD within a short evolutionary time frame, and the three changes were then fixed in all mouse species, potentially under positive selection. Therefore, we went on to examine whether these three amino acid changes caused any functional alteration of Cdx2.

**Fig 1.**
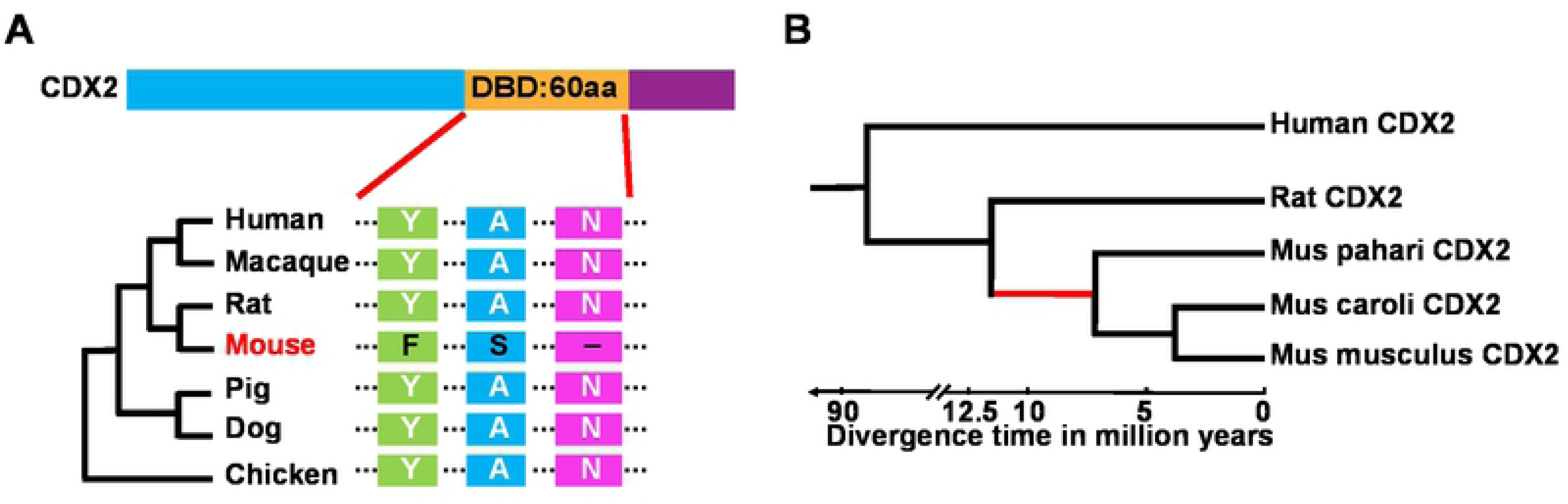
Three amino acid changes in the DNA binding domain of mCdx2. (A) Evolutionary comparison of Cdx2 from human to chicken indicated mouse had three specific amino acid changes in the DNA binding domain (DBD). (B) Phylogenetic analysis showed Cdx2 diverged between the *Rattus* and the *Mus* lineage about 5 to 12.5 million years ago.

### Cdx2 functions as a pioneer factor and induces trophoblast-like epigenomic and transcriptomic landscape

Cdx2 is a homeobox transcription factor essential for the development of the intestinal epithelium and the placenta [32, 36]. Serving as the first lineage specification marker [30–32, 37], previous studies have shown that ectopic overexpression of Cdx2 could efficiently promote the differentiation of ES cells to trophoblast stem cells [30, 31, 33]. Therefore, to investigate the function of Cdx2 during this process, we applied a doxycycline (DOX) inducible Tet-On system to induce Flag-tagged Cdx2 expression in mESCs and analyzed its function by measuring transcriptome and epigenome changes (Fig 2A). To check the suitability of this system, we first investigated the molecular function of mouse Cdx2 (mCdx2). As shown in Fig 2B, after successful induction of mCdx2 (Fig 2C, S2A and S2B Figs), typical round shape colonies of undifferentiated cells were disappearing, while differentiated epithelial-like cells with flat or square shape started to appear.

**Fig 2.**
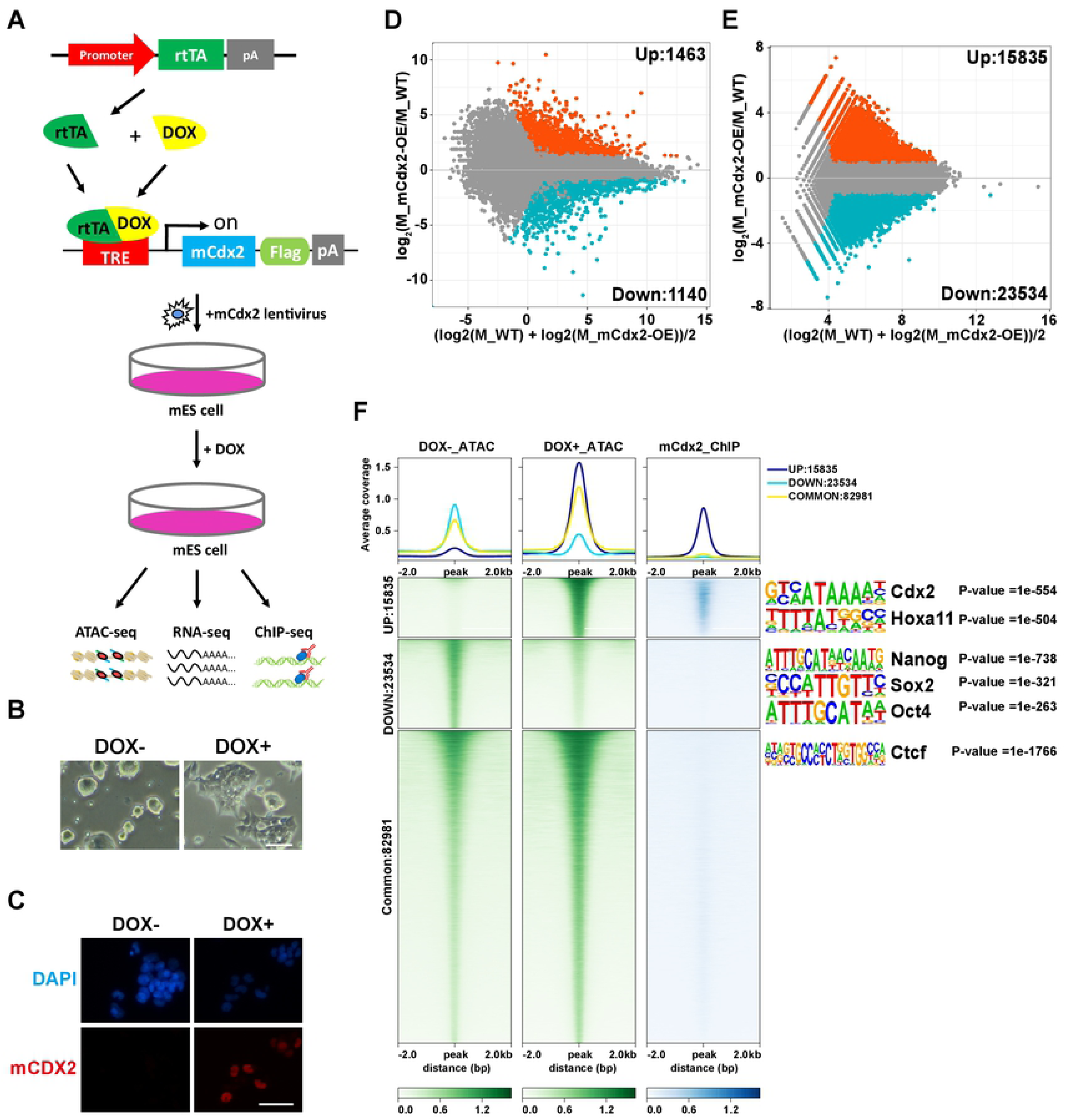
mCdx2 is an important regulator of stem cell differentiation. (A) Schematic diagram for the experimental design. Briefly, mESCs were transfected with mCdx2 lentivirus whose expression was controlled by DOX induction. After DOX treatment for 48h, cells were collected for RNA-seq, ATAC-seq and ChIP-seq analysis. (B) The cell morphology changed after DOX induction. (C) Immunofluorescence (IF) experiments confirmed the successful expression of mCdx2 using the antibody against CDX2. (D) The MA plot comparing expression profiles of representative ESC lines between DOX+ and DOX-conditions. Orange spots represented genes that showed higher than 2-fold expression (upregulated) and blue spots represented genes that show edlower than 2-fold expression (downregulated) in DOX+ condition than in DOX-condition. The number of up- and downregulated genes was also indicated. (E) The MA plot comparing the intensity of ATAC peaks between DOX+ and DOX-cells. The orange dots indicated the DOX+ specific peaks and the blue dots indicated the DOX-specific peaks. The number of upregulated and downregulated peaks were also indicated. (F) Aggregate plots showing the normalized read density of ATAC-peaks in mESCs with or without mCdx2 overexpression, along with mCdx2 binding sites. mCdx2 binding was directly associated with the openness of mCdx2-targeting ATAC-peaks. Corresponding motif enrichment in up, down and common ATAC-peaks were shown on the right.

We then compared the gene expression of the mESCs before and after DOX induction using RNA-seq (Fig 2A). As shown in Fig 2D, a total of 2603 genes exhibited significantly differential expression (|GFOLD|>1, TPM>3), with 1463 genes (56.2%) upregulated and 1140 (43.8%) genes downregulated after DOX induction. Notably, trophoblast stem cell (TSC) related markers such as Cdh2, Ascl2, Gata6 and Gata3 showed moderately increased expression while the expression of ESC markers such as Sox2 and Oct4 decreased, in consistence with the incipient trajectories of ES differentiation into TSCs (S2C Fig).

The massive differences in gene expression after mCdx2 induction suggested that dramatic differences in the epigenetic landscapes could be induced after mCdx2 overexpression (mCdx2-OE). To address this, ATAC-seq was carried out on cells with and without mCdx2-OE, respectively. A total of 39,369 peaks showed significantly different openness after mCdx2 induction (|log2 (fold change)|>1, total read count>=40, Fig 2E). To gain a deeper insight into the mCdx2 mediated epigenomic changes, we divided all the ATAC-peaks into three categories, namely UP (newly opened chromatin regions in mCdx2-OE cells), DOWN (chromatin accessibility lost in mCdx2-OE cells) and COMMON (regions without openness change) (Fig 2F), and then performed sequence motif analysis in these three categories separately. As shown in Fig 2F, UP regions were mostly enriched for DNA binding motifs of Cdx and Hox family transcription factors which are important for lineage specialization, while the DOWN motifs were mainly enriched for Oct4, Sox and Klf family genes which are known pluripotency regulators, and the motifs of the common peaks were enriched for house-keeping chromatin regulators, such as Ctcf (Fig 2F).

Last, to characterize mCdx2 binding sites on a genome-wide scale, we performed ChIP-seq in mCdx2-OE cells. A total of 93,591 peaks were identified, with 6.47%, 46.27% and 47.26% located at promoter, gene body, and distal intergenic regions, respectively (S2D Fig). Importantly, mCdx2 binding peaks were differentially distributed between the three categories of ATAC peaks, with predominant binding at the UP group (Fig 2F). Taken together, the results above revealed that mCdx2 likely acted as a pioneer transcription factor, instructing cell fate specification by binding, then opening chromatin and finally initiating the expression of important downstream target genes.

### Evolutionarily conserved function of mCdx2 and rCdx2 in transcriptional regulation

Then, to assess whether and how mouse Cdx2 might function differently as rat Cdx2 (rCdx2), we compared the effects of overexpressing mCdx2 and rCdx2. Like for mCdx2, we firstly established a Dox-inducible rCdx2 expression in mESCs. After 48h DOX treatment and successful rCdx2 induction, we observed similar morphological changes (S3A Fig) as with mCdx2 (S3B-D Figs). To explore the functional similarities and differences between rCdx2 and mCdx2 at the molecular level, we firstly carried out RNA-seq experiments on rCdx2-OE cells. At the transcriptome level, mCdx2-OE and rCdx2-OE cells showed high correlation (correlation = 0.992; p-value < 2.2 x 10^-16^, Fig 3A), suggesting their conserved effects on downstream gene expression. Then we conducted ChIP-seq to measure the rCdx2 DNA binding in rCdx2-OE mES cells. A total of 78,765 peaks were identified, of which 6.09%, 46.32% and 47.6% located at promoter, gene body, and distal intergenic regions, respectively (S3E Fig). The peak distribution pattern and the global binding signal showed nearly no differences between mCdx2 and rCdx2 (correlation=0.977, Figs 3B and 3C). As shown in Fig 3D, Hoxa family genes, well-known direct targets of Cdx2 exhibited the same DNA binding and gene expression change after mCdx2 and rCdx2 overexpression. Finally, we performed de novo motif analysis in mCdx2 and rCdx2 ChIP peaks, separately, and found an identical binding motif for mCdx2 and rCdx2, the same as previously reported for mCdx2 (Fig 3E). Collectively, these results indicated mCdx2 and rCdx2 are functionally conserved at the molecular level.

**Fig 3.**
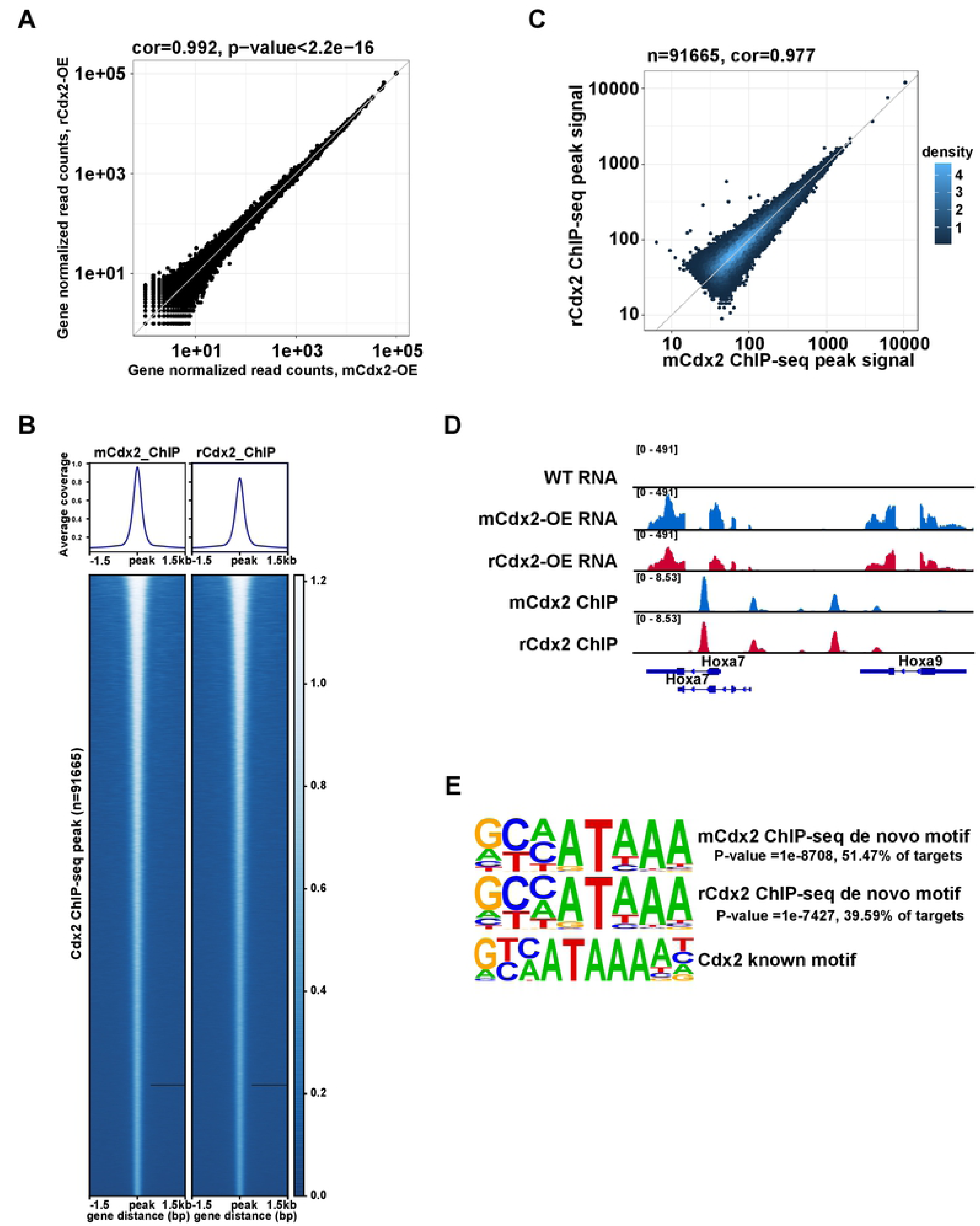
Functional comparison between mCdx2 and rCdx2. (A) The scatter plot comparing gene expression profile of mCdx2-OE and rCdx2-OE cells showed no significant differences. (B) Heat maps showed similar DNA binding patterns between mCdx2 and rCdx2. (C) Scatter-plot comparing ChIP-peak signal intensity showed high correlation of DNA binding affinity between mCdx2 and rCdx2. (D) mCdx2 and rCdx2 ChIP-Seq peaks at the Hoxa gene family region and associated gene expression pattern. mCdx2 signal were shown in blue and rCdx2 signal were shown in red. (E) Comparison of de novo binding motifs found in mCdx2 and rCdx2 ChIP-seq data, which were identical to the known Cdx2 binding motif.

### Mouse-specific amino acid changes in DBD, neither alone nor in combination have any effects on Cdx2 function

Based on the results above, the three amino acid changes in the DBD combined together did not induce any functional changes in our DOX-induced overexpression ES cells. Interestingly, whereas the three changes were observed after mouse-rat divergence, no other murine species were found to contain only one or two of the three changes. One possible evolutionary scenario was that one or two of the changes decreased DNA binding and were compensated by (a) later change(s) (i.e., the DBD found in mouse with all three amino acid changes in combination had the same binding affinity as that found in rat or human but changing only one or two of the amino acids could affect Cdx2 binding and function). To test this hypothesis, we constructed a panel of seven mutated plasmids consisting of all possible single-site changes, two-site changes and three-site changes in rCdx2 (Fig 4A and S4A Fig). Then we established stable cell lines transfected with these inducible mutated Cdx2s. After DOX treatment, all the cells tended to differentiate, manifested typical morphology changes (S4B Fig), and no obvious differences were found among all the mutated Cdx2 overexpressing cells. At the molecular level, we performed RNA-seq on cells overexpressing all different mutants (S4C Fig). As shown in Fig 4B, correlation analysis showed all the mutated Cdx2s had similar effects on the transcriptome compared to mCdx2 and rCdx2. Together, these results suggested no functional differences among the different mutants in our DOX-induced overexpression mESCs.

**Fig 4.**
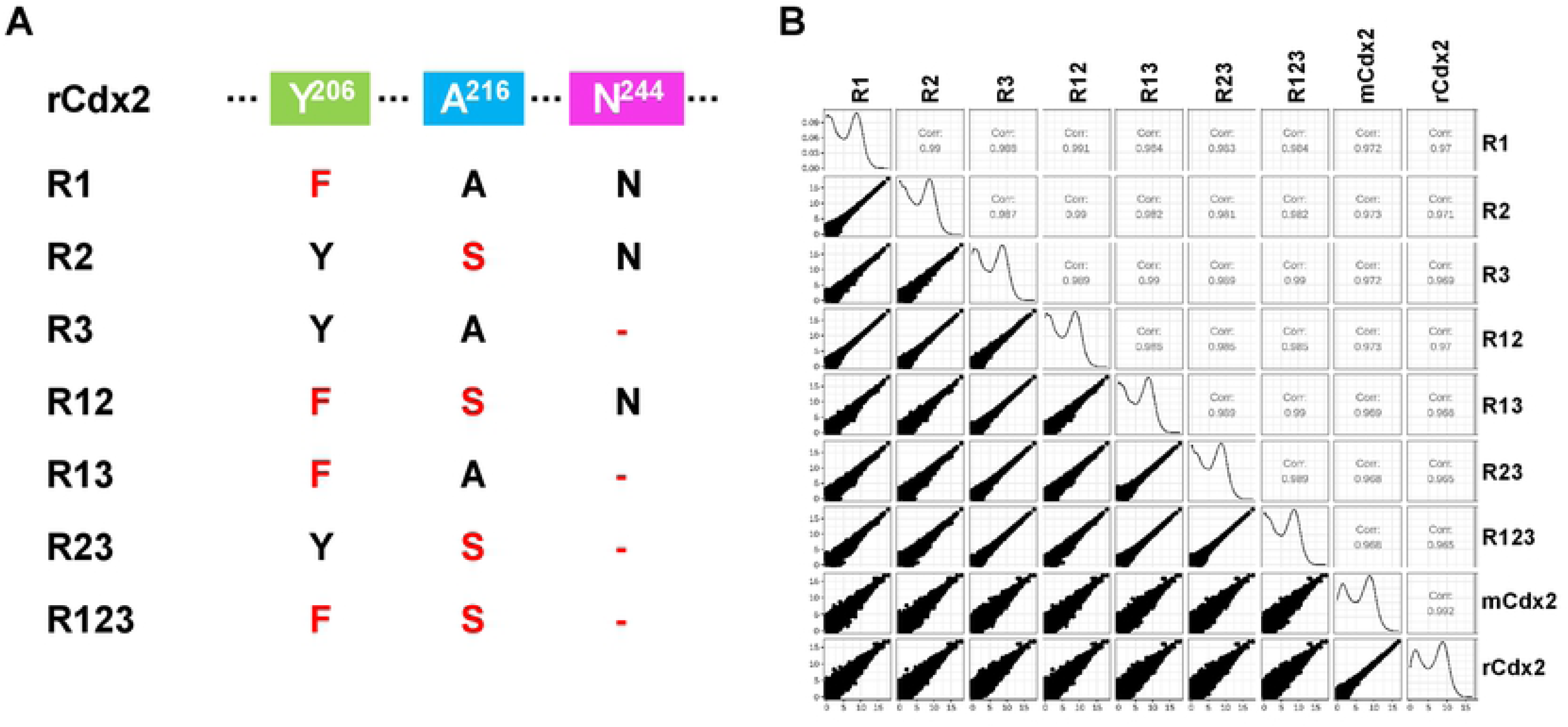
Conserved function of Cdx2 mutants. (A) The schematic representation of Cdx2 mutants. (B) The correlation analysis of RNA-seq results among mCdx2, rCdx2 and Cdx2 mutants.

### *Cis*-driven species-specific DNA binding of Cdx2

Transcriptional regulation of gene expression is mediated by both *cis* and *trans* components. The results shown above indicate that the amino acid differences in the Cdx2 DBD did not lead to divergent *trans*-regulatory effects between rat and mouse. We then turned to the *cis*-regulatory part of Cdx2-mediated regulation. One common strategy to study *cis*-effects is to compare the gene regulation between two alleles in F1 hybrid [3, 5, 12]. In F1 hybrids, both parental alleles are subject to the same *trans*-regulatory environments; thus, observed differences in allele-specific regulatory patterns should reflect only the impact of *cis* regulatory divergence. However, this approach cannot be conducted between mammalian species with long evolutionary distance, such as mouse and rat, due to reproductive isolation. To circumvent this limitation, we took advantage of the previously established RMES cells which contain a haploid mouse genome and a haploid rat genome [34]. Using this system, we then explored the *cis* divergence of Cdx2 mediated regulation between mouse and rat in an unbiased manner. Again, we established stable Cdx2-overexpressing RMES cell lines by transfecting Flag-tagged mCdx2 (mCdx2-OE RMESCs) and Flag-tagged rCdx2 (rCdx2-OE RMESCs), separately. Typical differentiation characteristics were observed after 48 hours DOX induction (S5A Fig). As before, RNA-seq and ChIP-seq were then conducted. Consistent with the observation in mESCs, mCdx2 and rCdx2 had similar DNA binding patterns in either mouse or rat genome, and induced similar gene expression changes on either rat or mouse allele in RMES cells (S5B-E Figs). In the following analysis, we combined the RNA-seq and ChIP-seq datasets from mCdx2-OE and rCdx2-OE RMESCs as experimental replicates.

Then we compared the Cdx2 binding sites between the mouse and rat genomes. In order to check how these binding sites evolved, we classified the binding sites determined by ChIP-seq based on whether the peaks in one species could be aligned to the other species and if yes, whether the aligned sites were also bound there (Fig 5A). The first two categories, including conserved and loss peaks represented the alignable sites, whereas the third one were those that could not be aligned to the other species. As shown in Fig 5B, conserved peaks, where the aligned binding sites were also bound, accounted for 23.9% (mouse to rat direction: 17,321 peaks) and 22.4% (rat to mouse direction: 17,317 peaks), respectively; loss peaks, referring to no binding in the orthologous regions, occupied 57.3% (mouse to rat direction: 41,430 peaks) and 58.5% (rat to mouse direction: 45,105 peaks), respectively; unaligned peaks constituted 18.8% (13,580 peaks, mouse to rat) and 19.1% (14,715 peaks, rat to mouse), respectively. The distributions of the three groups were similar between the binding sites at proximal and those at distal regions (S1 and S2 Tables).

**Fig 5.**
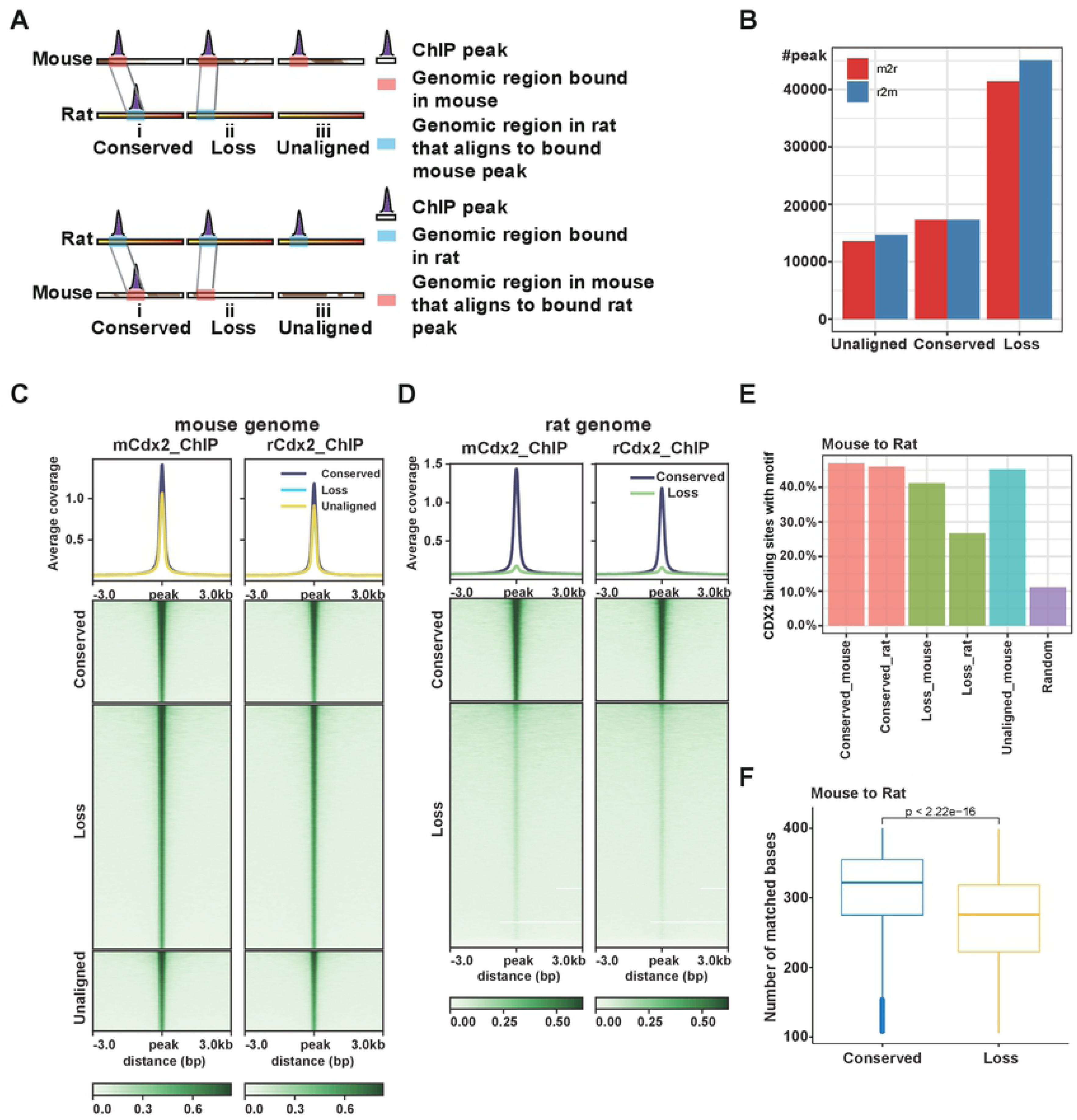
Species-specific binding of Cdx2. (A) ChIP-peaks were classified into three categoriest. In type (i) (conserved), the aligned regions were bound in both species; in type (ii) (loss), the orthologous sequence was found, but no binding was detected in the orthologous region; in type (iii) (unaligned), the aligned sequence was not present in the second species. (B) The counts of peak numbers in three categories, namely conserved, loss and unaligned. (C) The heatmap of signal intensity of three peak types at mouse genome and (D) after being aligned to the rat genome. (E) The percentage of ChIP-seq peaks with Cdx2 motif in three peak types from mouse to rat direction. (F) The distribution of number of matched bases between mouse and rat genome sequences in the conserved and loss peaks from mouse to rat direction.

Then we compared the signal intensities among these three peak classes. As expected, the binding affinities were the highest for conserved peaks and there were no significant differences between these conserved sites bound at the rat and mouse genomes (Figs 5C, 5D, S6A and S6B). Consistent with this, a gradual decrease in the frequency of Cdx2 binding motifs near binding sites was observed from conserved to loss peaks (Figs 5E, S6C Fig). To determine whether binding differences arise from sequence changes at potential binding sites, we checked the mutation rate among different peak categories. As expected, higher proportions of matched bases occurred in the conserved peaks relative to loss peaks (Fig 5F, S6D Fig). Therefore, the *cis*-driven DNA binding differences can be attributed to the sequence differences, particularly those affecting binding motifs.

### Cis-regulatory divergence of Cdx2 binding causes a fraction of species-specific gene expression differences

To further explore whether the divergent DNA binding could cause the species-biased gene expression, we compared Cdx2 induced transcriptional changes between mouse and rat alleles (Methods). As shown in Fig 6A, a total of 7,372 candidate genes were divided into five categories based on their expression changes after Cdx2 induction: type 0: no change, type 1: change only in the mouse allele, type 2: opposite changes between mouse and rat, type 3: change only in the rat allele, type 4: similar changes in both mouse and rat. A majority of genes (5592 genes, type 0) were not affected by Cdx2 induction for either the mouse or the rat allele; 496 genes (type 1) and 425 genes (type 3) with mouse-only and rat-only changes, respectively, reflected species-specific regulation; the type 2 category contained only 1 gene, indicating that opposite changes between mouse and rat after the same TF stimulus were extremely rare; a significant proportion of genes (858 genes, type 4) with similar expression changes after Cdx2 induction between mouse and rat suggested an evolutionarily conserved regulatory pattern (Fig 6B). Based on the GO analysis, we found that type 4 genes were highly enriched in functions related to “organism development and cell differentiation, consistent with the known function of Cdx2 in early development (Fig 6C). In addition, the magnitude of gene expression changes was highest for type 4 genes, further suggesting their important functions (Fig 6D).

**Fig 6.**
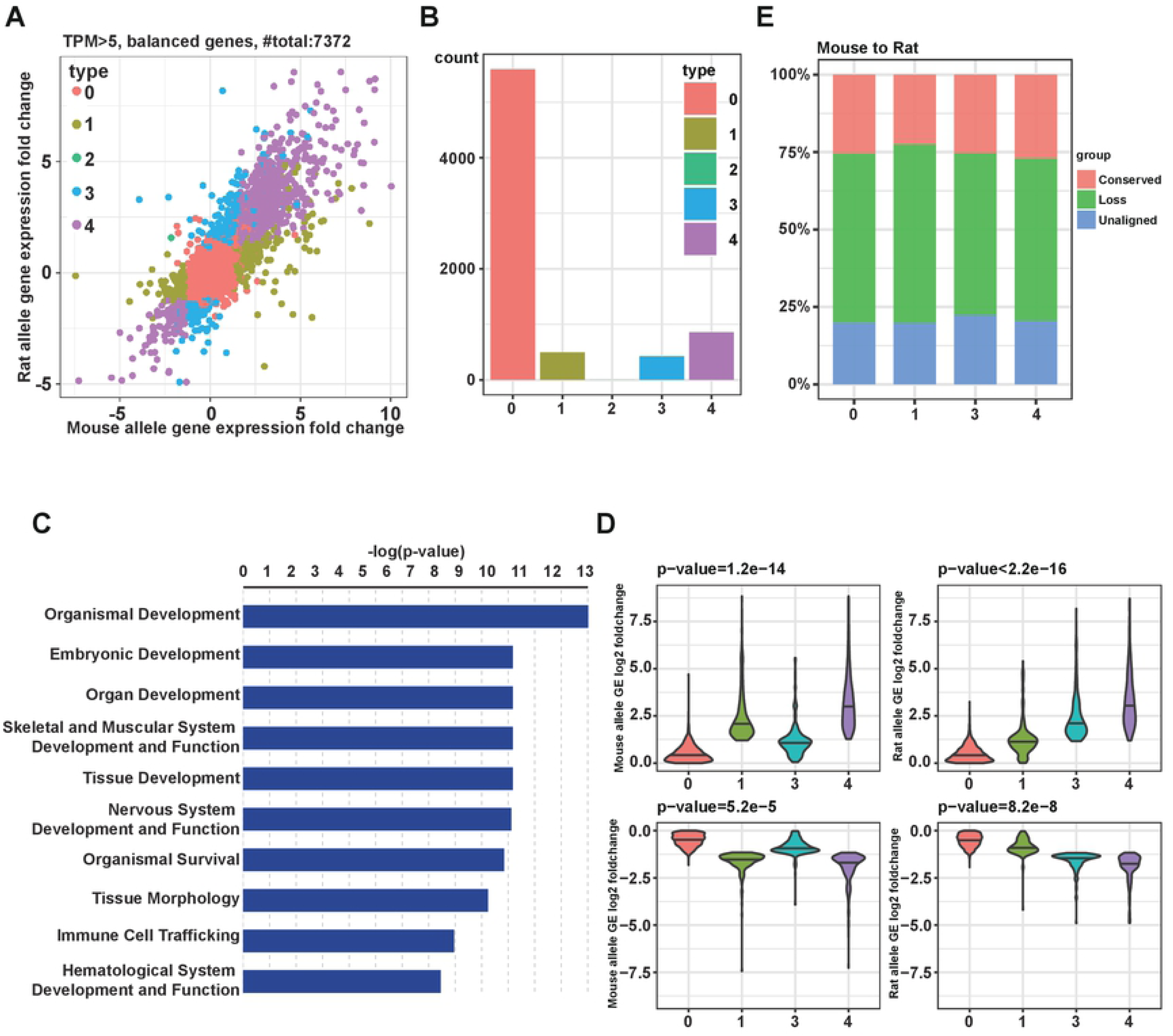
The relationship of differential DNA binding and gene expression. (A) The gene expression changes after Cdx2 induction for mouse allele and rat allele respectively. Genes were divided into five groups based on the change direction between mouse allele and rat allele. For the detail, see the main text. (B) The numbers of genes in the five groups (0, 1, 2, 3, 4). (C) Gene ontology (GO) enrichment analysis using Ingenuity Pathway Analysis (IPA) software (QIAGEN). Genes of type 4 were enriched for functions related to “organism development and cell differentiation”. (D) The magnitude of gene expression changes of type 0,1,3,4 genes. (E) The percentage of different kinds of peaks in four groups (0, 1, 3, 4) of genes from mouse to rat direction.

Finally, we analyzed the relationship between differential binding and gene expression. For this purpose, we checked the distribution of peaks located in different groups of genes. As expected, the proportion of conserved peaks in type 4 was moderately higher than in that of the other types (Figs 6E, S6E Fig). Similarly, the relatively higher percentage of loss peaks in type 1 and unaligned peaks in type 3 suggested the role of these species-specific binding in species-specific regulation. Nevertheless, among type 4 genes with conserved expression changes, more than 70% had no Cdx2 binding sites conserved between the two species. Therefore, at the *cis* regulatory level, whereas the Cdx2 binding was largely divergent between mouse and rat, the transcriptional effect induced by Cdx2 was conserved to a much larger extent.

## Discussion

In this study, we systematically sought species specific mutations in the DBD of TFs and identified Cdx2 as an apparent candidate for adaptive evolution in the mouse lineage with three specific amino acid changes exclusively present in mCdx2 DBD. Given that Cdx2 plays an important role in embryonic development and stem cell differentiation [31, 32, 37], we investigated its function in mESC differentiation. In line with previous studies [30], induction of mCdx2 caused dramatic transcriptomic and epigenetic changes, confirming the crucial role of mCdx2 in cell differentiation and lineage specification. Next, we chose rCdx2 as an example to compare its function with mCdx2. The results indicated that mCdx2 and rCdx2 were functionally conserved at the molecular level. Furthermore, similar conclusions could be drawn when examining the effects of mouse-specific amino acid changes alone or in combination. These data together suggested that the regulatory function of Cdx2 was evolutionarily conserved and was unaffected by any of the derived mouse-specific amino acid changes, which contradicted our initial expectation. It is of course possible that these mouse-specific mutations are the result of neutral drift rather than positive selection, and the amino acid changes do not affect the binding affinity and function of Cdx2 in any significant way. Therefore it would be interesting to see whether any extant murine species carries Cdx2 with only one or two of the mouse-specific amino acid changes, when genome sequencing efforts progress. Alternatively, it is also possible that the exogenous overexpression of Cdx2 might mask the subtle species-specific effect which might be present at normal physiological level, in other words, some false negatives for binding differences between mCdx2 and rCdx2 could be due to high expression of Cdx2. Furthermore, the function of Cdx2 may be context dependent, meaning these mutations may lead to divergent functions at a later stage during development or in other tissues where Cdx2 is expressed (e.g., adult intestine).

We then analyzed *cis*-regulatory divergence on Cdx2 mediated regulation. As to transcriptional effects, Cdx2 mediated regulation appeared to be largely conserved between mouse and rat. Among genes showing Cdx2 regulation in mouse or rat alleles, 48.2% genes displayed a conserved regulatory pattern between the two alleles. In addition, the induced change is also of higher magnitude for those genes with conserved regulation. Furthermore, these genes were enriched in functions related to development and differentiation, consistent with the established function of Cdx2 in early development. In contrast, Cdx2 binding sites are largely divergent between the mouse and rat allele, with only about 23% of peaks being conserved. Several possibilities could account for this apparent discrepancy. First, Cdx2 mostly binds to distal regions, therefore its target genes are difficult to define. Furthermore, not all TF binding events could lead to transcriptional output. On the one hand, many binding events could be redundant in regulating the expression of their target genes. On the other hand, in some cases, binding by co-factor(s) would be required to exert transcriptional impact. In addition, many observed binding events would be non-functional biological noise [38], particularly with the ectopic overexpression of TF, as we did here. Last not the least, as observed in a previous study, TF binding locations can frequently change, even though gene expression is conserved between human and mouse, suggesting a high functional plasticity and fast evolvability of TF binding [39, 40].

Finally, we also explored the potential mechanisms underlying the differences in Cdx2 binding between the two species. The conserved peaks had the highest proportions of matched bases, suggesting the association between genetic sequence variation and TF binding differences between alleles. Nevertheless, the loss of binding in spite of high sequence/motif conservation was also observed. In addition, we could find many cases with conserved binding in spite of loss of the Cdx2 motif in one allele. Both of these observations implied a role of other TFs in Cdx2 binding at these sites. These cofactors in Cdx2 mediated gene regulation await future analysis.

## Materials and Methods

### Cell culture

The mESC and RMES cells were cultured as previously reported [34]. In brief, cells were maintained in Neuralbasal (Invitrogen, cat. no. 21103-049)-DMEM/F12 (Invitrogen, cat. no. 11330-032) based medium supplemented with N2 (Invitrogen, cat. no. 17502048), B27 (Invitrogen, cat. no. 17504-044), PD0325901 (Selleck, cat. no. s1036), Chir99021 (Selleck, cat. no. s1263) and mLIF (Millipore, cat. no. ESG1107) at 5% CO_2_ and 37°C.

HEK 293T cells were maintained in DMEM medium (Gibco, USA) supplemented with 10% FBS (Gibco, USA) and 1% penicillin/streptomycin (Thermo) at 5% CO2 and 37°C.

### Plasmid vector construction

mCdx2 and rCdx2 overexpression (OE) vectors were constructed by replacing APEX2 with mCdx2 or rCdx2 in the RAR3G - APEX2 - FLAG plasmid [41] based on the homologous recombination technology (Vazyme C112). The mutated Cdx2 plasmids were modified from the rCdx2 plasmid based on ClonExpress rapid cloning technology according to the manufacturer’s instructions (Vazyme C215). In brief, the fragments were amplified respectively according to the mutation sites followed by DpnI digestion to remove the methylated template plasmid, and then these mutated fragments were recombined with the vector. Recombination product was transformed directly to complete the multiple base site-directed mutagenesis.

### Lentivirus package and transfection

All the target plasmids were co-transfected into the HEK 293T cells with the helper plasmids psPAX2 and pMD2.G. 72 hours later, we collected the liquid supernatant to filter and concentrate the final virus. Then the virus was added to the cultured cells to get the stable cell lines. mESC transfected with mCdx2 and rCdx2 were stimulated with DOX (200 ng/mL) for 48 h to induce the expression of Cdx2.

### Immunofluorescence assay

For immunofluorescent staining, cells were fixed in 4% formaldehyde at 4°C overnight. After washing with cold PBS, cells were permeabilized with 0.1% Triton X-100 in PBS for 15 min and incubated with block buffer (3% BSA) for 30 min before being probed with primary antibodies. Then, cells were stained with CDX2 antibodies (anti-rabbit, 1:300) and FLAG antibodies (anti-mouse,1:300) at 4°C overnight, followed by secondary antibody for 30 min at room temperature. Nuclei were stained with DAPI (1:5000). The images were obtained by fluorescent microscopy (Nikon, Japan), and processed by Adobe Photoshop 2019.

### RNA extraction from cultured cells

Total RNA were extracted from cells using TRIzol reagent according to the manufacturer’s protocol (Life Technologies). The integrity of purified total RNA was estimated by Agilent Bioanalyzer using RNA Nano kit (Agilent Technologies) before subsequent experiments. Total RNA with an RNA integrity number (RIN) above 9.0 was used for mRNA-seq.

### mRNA-seq

Truseq Stranded mRNA sequencing libraries were prepared with 2 μg total RNA according to the manufacturer’s protocol (Yeasen 12300ES24). The libraries were sequenced in 2 × 150 nt manner on HiSeq Xten platform (Illumina).

### ATAC-seq library preparation and sequencing

For ATAC library construction, cells were washed and then lysed in 50 μL lysis buffer (10 mM Tris-HCl (pH 7.4), 10 mM NaCl, 3 mM MgCl_2_, 0.1% NP-40, 0.1% Tween-20, and 0.01% digitonin) for 3 min on ice. Immediately after lysis, samples were then incubated with the Tn5 transposase and tagmentation buffer at 37°C for 30 min (Vazyme Biotech, TD501). PCR was then performed to amplify the library for 12 cycles using the following PCR cycles: 72°C for 3 min; 98°C for 30 s followed by thermocycling at 98°C for 15 s, 60°C for 30 s and 72°C for 40s, and finally 5 min at 72°C. After PCR, libraries were purified with 1.2X DNA clean beads (Vazyme, N411). The libraries were sequenced in 2 × 150 nt manner on HiSeq Xten platform (Illumina).

### Chromatin immunoprecipitation followed by sequencing (ChIP-seq)

The ChIP assay was performed according to the standard protocol provided by SimpleChIP Plus Sonication Chromatin IP Kit (CST, 56383) with minor modifications. Briefly, 10^7^ cells were fixed with 1% formaldehyde for 10 minutes. Sonication was then carried out at the Bioruptor pico (Diagenode) by applying 10 cycles of 30 seconds ON and 30 seconds OFF to obtain chromatin fragments of approximately 100-500 bp. ChIP was performed with the anti-Flag M2 antibody (F1804, Sigma). ChIP DNA was cleaned up using the ChIP DNA Clean & Concentrator kit (Zymo, D5205). ChIP-seq libraries were prepared using standard protocol provided by VAHTSTM Universal DNA Library Prep Kit for Illumina® V3 (Vazyme, ND607). The libraries were sequenced in 2 × 150 nt manner on HiSeq Xten platform (Illumina).

### Multiple sequence alignment

Protein sequences of CDX2 of 56 species were extracted from UniProt (https://www.uniprot.org/). Multiple sequence alignment of CDX2 protein sequences were performed by MUSCLE [42] with default parameters, and visualized using MView [43].

### RNA-seq data analysis

STAR (v2.7.0d) [44] was used to align the RNA-seq reads to mouse genome (mm10) with gene annotation. The gene annotation gtf file was obtained from Ensemble v92 and pseudogenes were removed. FeatureCounts (v1.5.3) [45] was used to count the read number of reads mapped with each gene (with parameters -T 12 -s 2 -p -g gene_name). Gfold [46] was used to identify the differential expressed genes (DEG) if there was only one replicate. When there were at least two replicates, DESeq2 [47] was used to determine the DEG.

Homologous gene relationship between mouse and rat was downloaded from Ensembl BioMart (https://www.ensembl.org/biomart/martview).For RMES cells, reads were aligned to both mm10 and rn6 reference respectively by STAR, and assigned to either mm10 or rn6 according to the alignment edit distance to the reference.

### ATAC-seq data analysis

Fastp (v0.19.5) [48] was first used to trim and remove low quality reads and adapter sequences (-a CTGTCTCTTATA --detect_adapter_for_pe -w 12 --length_required 20 -q 30). Reads were aligned to the mouse reference genome (mm10) using Bowtie2 [49] v2.3.4.3 with parameters (-p 12 -X 2000). Reads mapped to the mitochondrial genome and low mapping quality reads (MAPQ < 20) were filtered out using custom scripts. Sambamba [50] (v0.7.0) was then used to sort the reads and remove duplicates. Reads from both mESC with and without Cdx2-OE samples were merged as input for MACS2 [51] to call peaks (-g mm --keep-dup all -q 0.05 --slocal 10000 -- nomodel --nolambda). To control the false positives, high confident ATAC peaks (-log10(q-value) >4) were used in the following analyses. FeatureCounts v1.5.3 was used to count the read number within each ATAC peak in each sample. Differential peaks between mESC with and without Cdx2-OE were identified by comparing the normalized read counts (|Log2 (Fold change)| > 1, total normalized read count no less than 40). The remaining peaks were referred to as common peaks. HOMER [52] was used to perform motif enrichment analysis on the ATAC peaks. Deeptools [53] was used to plot the heatmap of peak signal across different regions.

### ChIP-seq data analysis

2×150 bp paired-end reads were first trimmed to remove adapter sequences using cutadapt[54] v2.6 (-a AGATCGGAAGAG-A AGATCGGAAGAG -m 20 -q 30). Reads were aligned to mouse reference genome (mm10) using Bowtie2 v2.3.4.3 with parameters (-X 2000). Reads mapped to mitochondrial genome and low mapping quality reads (MAPQ < 20) were filtered out using custom script. Picard (v2.6.0, http://broadinstitute.github.io/picard/) was then used to sort the reads and remove duplicates.

To systematically compare mCDX2 ChIP-seq and rCDX2 ChIP-seq peak signals, reads from both mCDX2 and rCDX2 ChIP-seq samples were merged as input for MACS2 to call peaks (-g mm --keep-dup all -q 0.05 -B). To control the false positives, 91665 high confidence ChIP-seq peaks (-log10 (q-value) >4) were used in the following analysis. FeatureCounts v1.6.0 was used to count the read number within each peak in each sample. R package ChIPseeker [55] was used to annotate the peaks. Fimo with default parameters was used to determine the CDX2 motif occurrence within the ChIP peaks.

For RMES cells, reads were aligned to both mm10 and rn6 reference respectively by Bowtie2, and assigned to either mm10 or rn6 according to the alignment edit distance to the reference. The subsequent analysis was the same as the single species data analysis above.

### Cross species peak analysis

All peaks were centered at peak summit and extend 200 bp on both sides to control length. DNA sequences of the peaks were extracted by bedtools [56] from the genome. pblat [57], a multithread version of blat was used to align the peak sequences to the other genome with parameters (-t=dna -q=dna-minIdentity=70 -minScore=100 -ooc x.11.ooc). pslReps (-singleHit -minAli=0.7 -nohead) was used to generate genome-wide best alignments from the output of pblat. Multiple-mapped peaks were excluded in this analysis. Peak classification criteria were defined as in a previous study [39]. Briefly, peaks were classified according to whether corresponding aligned regions exist in the second species and whether these aligned regions were bound with a peak. If the peak sequence was not alignable to the second species, it was classified as unaligned. If the aligned region overlaps more than 50% of a ChIP peak, it was classified as conserved. The other was classified as loss.

### Linking genes to Cdx2 binding

To determine the Cdx2 binding composition of the homologous genes, 60 kb region centered at gene start sites were used to intersect with ChIP peaks. The proportion of different group of peaks for each type of genes was calculated. Because Type 2 contains 1 gene, this type was excluded for this analysis.

### Gene ontology (GO) enrichment analysis

GO enrichment analysis was carried by subjecting gene sets to the Ingenuity Pathway Analysis (IPA) (QIAGEN) and the “Core Analysis” of “Expression Analysis” was performed. In the “Diseases & Function” module, the analysis was restricted to “Physiological System Development and Function”.

## Acknowledgments

This work was supported by Shenzhen Key Laboratory of Gene Regulation and Systems Biology, Shenzhen-Hong Kong Institute of Brain Science-Shenzhen Fundamental Research Institutions (Grant No. 2021SHIBS0002), Shenzhen Science and Technology Program (Grant No. KQTD20180411143432337 and JCYJ20180504165804015), the National Natural Science Foundation of China (Grant No. 31771443, 31621004, 31900431, 31900462, 31970821), and National Key Research and Development Program (2019YFA0110800 and 2019YFA0903802). The authors acknowledge the Center for Computational Science and Engineering of SUSTech for the support on computational resource and acknowledge the SUSTech Core Research Facilities for technical supports.

## References

1. Hinman V, Cary G. The evolution of gene regulation. eLife. 2017;6:e27291. doi: 10.7554/eLife.27291.

2. King MC, Wilson AC. Evolution at two levels in humans and chimpanzees. Science. 1975;188(4184):107–16. Epub 1975/04/11. doi: 10.1126/science.1090005. PubMed PMID: 1090005.

3. Wittkopp PJ, Haerum BK, Clark AG. Evolutionary changes in cis and trans gene regulation. Nature. 2004;430(6995):85–8. doi: 10.1038/nature02698. PubMed PMID: WOS:000222356800048.

4. Signor SA, Nuzhdin SV. The Evolution of Gene Expression in cis and trans. Trends Genet. 2018;34(7):532–44. doi: 10.1016/j.tig.2018.03.007. PubMed PMID: WOS:000435027200005.

5. Goncalves A, Leigh-Brown S, Thybert D, Stefflova K, Turro E, Flicek P, et al. Extensive compensatory cis-trans regulation in the evolution of mouse gene expression. Genome Res. 2012;22(12):2376–84. doi: 10.1101/gr.142281.112. PubMed PMID: WOS:000311895500006.

6. Meiklejohn CD, Coolon JD, Hartl DL, Wittkopp PJ. The roles of cis- and trans-regulation in the evolution of regulatory incompatibilities and sexually dimorphic gene expression. Genome Res. 2014;24(1):84–95. doi: 10.1101/gr.156414.113. PubMed PMID: WOS:000329163500008.

7. Wray GA, Hahn MW, Abouheif E, Balhoff JP, Pizer M, Rockman MV, et al. The evolution of transcriptional regulation in eukaryotes. Mol Biol Evol. 2003;20(9):1377–419. doi: 10.1093/molbev/msg140. PubMed PMID: WOS:000185175500001.

8. Yvert G, Brem RB, Whittle J, Akey JM, Foss E, Smith EN, et al. Trans-acting regulatory variation in Saccharomyces cerevisiae and the role of transcription factors. Nat Genet. 2003;35(1):57–64. Epub 2003/08/05. doi: 10.1038/ng1222. PubMed PMID: 12897782.

9. Pickrell JK, Marioni JC, Pai AA, Degner JF, Engelhardt BE, Nkadori E, et al. Understanding mechanisms underlying human gene expression variation with RNA sequencing. Nature. 2010;464(7289):768–72. Epub 2010/03/12. doi: 10.1038/nature08872. PubMed PMID: 20220758; PubMed Central PMCID: PMCPMC3089435.

10. Gao Q, Sun W, Ballegeer M, Libert C, Chen W. Predominant contribution of cis-regulatory divergence in the evolution of mouse alternative splicing. Mol Syst Biol. 2015;11(7):816. Epub 2015/07/03. doi: 10.15252/msb.20145970. PubMed PMID: 26134616; PubMed Central PMCID: PMCPMC4547845.

11. Hou J, Wang X, McShane E, Zauber H, Sun W, Selbach M, et al. Extensive allele-specific translational regulation in hybrid mice. Mol Syst Biol. 2015;11(8):825. Epub 2015/08/09. doi: 10.15252/msb.156240. PubMed PMID: 26253569; PubMed Central PMCID: PMCPMC4562498.

12. Wittkopp PJ, Haerum BK, Clark AG. Regulatory changes underlying expression differences within and between Drosophila species. Nat Genet. 2008;40(3):346–50. doi: 10.1038/ng.77. PubMed PMID: WOS:000253548400027.

13. Wong ES, Schmitt BM, Kazachenka A, Thybert D, Redmond A, Connor F, et al. Interplay of cis and trans mechanisms driving transcription factor binding and gene expression evolution. Nat Commun. 2017;8. doi: ARTN 1092 10.1038/s41467-017-01037-x. PubMed PMID: WOS:000413404800003.

14. Bell GD, Kane NC, Rieseberg LH, Adams KL. RNA-seq analysis of allele-specific expression, hybrid effects, and regulatory divergence in hybrids compared with their parents from natural populations. Genome biology and evolution. 2013;5(7):1309–23. Epub 2013/05/17. doi: 10.1093/gbe/evt072. PubMed PMID: 23677938; PubMed Central PMCID: PMCPMC3730339.

15. Chen J, Nolte V, Schlötterer C. Temperature stress mediates decanalization and dominance of gene expression in Drosophila melanogaster. PLoS Genet. 2015;11(2):e1004883. Epub 2015/02/27. doi: 10.1371/journal.pgen.1004883. PubMed PMID: 25719753; PubMed Central PMCID: PMCPMC4342254.

16. Coolon JD, McManus CJ, Stevenson KR, Graveley BR, Wittkopp PJ. Tempo and mode of regulatory evolution in Drosophila. Genome Res. 2014;24(5):797–808. Epub 2014/02/26. doi: 10.1101/gr.163014.113. PubMed PMID: 24567308; PubMed Central PMCID: PMCPMC4009609.

17. Emerson JJ, Hsieh LC, Sung HM, Wang TY, Huang CJ, Lu HH, et al. Natural selection on cis and trans regulation in yeasts. Genome Res. 2010;20(6):826–36. Epub 2010/05/07. doi: 10.1101/gr.101576.109. PubMed PMID: 20445163; PubMed Central PMCID: PMCPMC2877579.

18. Gruber JD, Vogel K, Kalay G, Wittkopp PJ. Contrasting properties of gene-specific regulatory, coding, and copy number mutations in Saccharomyces cerevisiae: frequency, effects, and dominance. PLoS Genet. 2012;8(2):e1002497. Epub 2012/02/22. doi: 10.1371/journal.pgen.1002497. PubMed PMID: 22346762; PubMed Central PMCID: PMCPMC3276545.

19. Metzger BP, Duveau F, Yuan DC, Tryban S, Yang B, Wittkopp PJ. Contrasting Frequencies and Effects of cis- and trans-Regulatory Mutations Affecting Gene Expression. Molecular biology and evolution. 2016;33(5):1131–46. Epub 2016/01/20. doi: 10.1093/molbev/msw011. PubMed PMID: 26782996; PubMed Central PMCID: PMCPMC4909133.

20. Schaefke B, Emerson JJ, Wang TY, Lu MY, Hsieh LC, Li WH. Inheritance of gene expression level and selective constraints on trans- and cis-regulatory changes in yeast. Molecular biology and evolution. 2013;30(9):2121–33. Epub 2013/06/25. doi: 10.1093/molbev/mst114. PubMed PMID: 23793114.

21. Sung HM, Wang TY, Wang D, Huang YS, Wu JP, Tsai HK, et al. Roles of trans and cis variation in yeast intraspecies evolution of gene expression. Molecular biology and evolution. 2009;26(11):2533–8. Epub 2009/08/04. doi: 10.1093/molbev/msp171. PubMed PMID: 19648464; PubMed Central PMCID: PMCPMC2767097.

22. Suvorov A, Nolte V, Pandey RV, Franssen SU, Futschik A, Schlötterer C. Intra-specific regulatory variation in Drosophila pseudoobscura. PLoS One. 2013;8(12):e83547. Epub 2014/01/05. doi: 10.1371/journal.pone.0083547. PubMed PMID: 24386226; PubMed Central PMCID: PMCPMC3873948.

23. Wang D, Sung HM, Wang TY, Huang CJ, Yang P, Chang T, et al. Expression evolution in yeast genes of single-input modules is mainly due to changes in trans-acting factors. Genome Res. 2007;17(8):1161–9. Epub 2007/07/07. doi: 10.1101/gr.6328907. PubMed PMID: 17615293; PubMed Central PMCID: PMCPMC1933509.

24. Metzger BPH, Wittkopp PJ, Coolon JD. Evolutionary Dynamics of Regulatory Changes Underlying Gene Expression Divergence among Saccharomyces Species. Genome Biol Evol. 2017;9(4):843–54. doi: 10.1093/gbe/evx035. PubMed PMID: WOS:000406755800003.

25. Crowley JJ, Zhabotynsky V, Sun W, Huang S, Pakatci IK, Kim Y, et al. Analyses of allele-specific gene expression in highly divergent mouse crosses identifies pervasive allelic imbalance. Nat Genet. 2015;47(4):353–60. Epub 2015/03/03. doi: 10.1038/ng.3222. PubMed PMID: 25730764; PubMed Central PMCID: PMCPMC4380817.

26. Villar D, Flicek P, Odom DT. Evolution of transcription factor binding in metazoans - mechanisms and functional implications. Nat Rev Genet. 2014;15(4):221–33. doi: 10.1038/nrg3481. PubMed PMID: WOS:000333191900009.

27. Nitta KR, Jolma A, Yin Y, Morgunova E, Kivioja T, Akhtar J, et al. Conservation of transcription factor binding specificities across 600 million years of bilateria evolution. Elife. 2015;4. Epub 2015/03/18. doi: 10.7554/eLife.04837. PubMed PMID: 25779349; PubMed Central PMCID: PMCPMC4362205.

28. Slodkowicz G, Goldman N. Integrated structural and evolutionary analysis reveals common mechanisms underlying adaptive evolution in mammals. Proc Natl Acad Sci U S A. 2020;117(11):5977–86. Epub 2020/03/04. doi: 10.1073/pnas.1916786117. PubMed PMID: 32123117; PubMed Central PMCID: PMCPMC7084095.

29. Olson-Manning CF, Wagner MR, Mitchell-Olds T. Adaptive evolution: evaluating empirical support for theoretical predictions. Nature reviews Genetics. 2012;13(12):867–77. Epub 2012/11/17. doi: 10.1038/nrg3322. PubMed PMID: 23154809; PubMed Central PMCID: PMCPMC3748133.

30. Nishiyama A, Xin L, Sharov AA, Thomas M, Mowrer G, Meyers E, et al. Uncovering early response of gene regulatory networks in ESCs by systematic induction of transcription factors. Cell Stem Cell. 2009;5(4):420–33. Epub 2009/10/03. doi: 10.1016/j.stem.2009.07.012. PubMed PMID: 19796622; PubMed Central PMCID: PMCPMC2770715.

31. Jana D, Kale HT, Shekar PC. Generation of Cdx2-mCherry knock-in murine ES cell line to model trophectoderm and intestinal lineage differentiation. Stem cell research. 2019;39:101521. Epub 2019/08/11. doi: 10.1016/j.scr.2019.101521. PubMed PMID: 31400702.

32. Amin S, Neijts R, Simmini S, van Rooijen C, Tan SC, Kester L, et al. Cdx and T Brachyury Co-activate Growth Signaling in the Embryonic Axial Progenitor Niche. Cell Rep. 2016;17(12):3165–77. Epub 2016/12/24. doi: 10.1016/j.celrep.2016.11.069. PubMed PMID: 28009287.

33. Peng K, Li X, Wu C, Wang Y, Yu J, Zhang J, et al. Derivation of Haploid Trophoblast Stem Cells via Conversion In Vitro. iScience. 2019;11:508–18. Epub 2019/01/13. doi: 10.1016/j.isci.2018.12.014. PubMed PMID: 30635215; PubMed Central PMCID: PMCPMC6354440.

34. Li X, Cui XL, Wang JQ, Wang YK, Li YF, Wang LY, et al. Generation and Application of Mouse-Rat Allodiploid Embryonic Stem Cells. Cell. 2016;164(1-2):279–92. Epub 2016/01/16. doi: 10.1016/j.cell.2015.11.035. PubMed PMID: 26771496.

35. Thybert D, Roller M, Navarro FCP, Fiddes I, Streeter I, Feig C, et al. Repeat associated mechanisms of genome evolution and function revealed by the Mus caroli and Mus pahari genomes. Genome Res. 2018;28(4):448–59. Epub 2018/03/23. doi: 10.1101/gr.234096.117. PubMed PMID: 29563166; PubMed Central PMCID: PMCPMC5880236.

36. Kumar N, Tsai YH, Chen L, Zhou A, Banerjee KK, Saxena M, et al. The lineage-specific transcription factor CDX2 navigates dynamic chromatin to control distinct stages of intestine development. Development. 2019;146(5). Epub 2019/02/13. doi: 10.1242/dev.172189. PubMed PMID: 30745430; PubMed Central PMCID: PMCPMC6432663.

37. Cui T, Jiang L, Li T, Teng F, Feng G, Wang X, et al. Derivation of Mouse Haploid Trophoblast Stem Cells. Cell Rep. 2019;26(2):407–14.e5. Epub 2019/01/10. doi: 10.1016/j.celrep.2018.12.067. PubMed PMID: 30625323.

38. Spivakov M. Spurious transcription factor binding: Non-functional or genetically redundant? Bioessays. 2014;36(8):798–806. PubMed PMID: WOS:000339551700012.

39. Odom DT, Dowell RD, Jacobsen ES, Gordon W, Danford TW, MacIsaac KD, et al. Tissue-specific transcriptional regulation has diverged significantly between human and mouse. Nat Genet. 2007;39(6):730–2. doi: 10.1038/ng2047. PubMed PMID: WOS:000246859100016.

40. Dermitzakis ET, Clark AG. Evolution of transcription factor binding sites in mammalian gene regulatory regions: Conservation and turnover. Mol Biol Evol. 2002;19(7):1114–21. doi: DOI 10.1093/oxfordjournals.molbev.a004169. PubMed PMID: WOS:000176697400012.

41. Ke M, Liu J, Chen W, Chen L, Gao W, Qin Y, et al. Integrated and Quantitative Proteomic Approach for Charting Temporal and Endogenous Protein Complexes. Analytical chemistry. 2018;90(21):12574–83. Epub 2018/10/04. doi: 10.1021/acs.analchem.8b02667. PubMed PMID: 30280895.

42. Edgar RC. MUSCLE: a multiple sequence alignment method with reduced time and space complexity. BMC bioinformatics. 2004;5:113. Epub 2004/08/21. doi: 10.1186/1471-2105-5-113. PubMed PMID: 15318951; PubMed Central PMCID: PMCPMC517706.

43. Brown NP, Leroy C, Sander C. MView: a web-compatible database search or multiple alignment viewer. Bioinformatics. 1998;14(4):380–1. Epub 1998/06/20. doi: 10.1093/bioinformatics/14.4.380. PubMed PMID: 9632837.

44. Dobin A, Davis CA, Schlesinger F, Drenkow J, Zaleski C, Jha S, et al. STAR: ultrafast universal RNA-seq aligner. Bioinformatics. 2013;29(1):15–21. Epub 2012/10/30. doi: 10.1093/bioinformatics/bts635. PubMed PMID: 23104886; PubMed Central PMCID: PMCPMC3530905.

45. Liao Y, Smyth GK, Shi W. featureCounts: an efficient general purpose program for assigning sequence reads to genomic features. Bioinformatics. 2014;30(7):923–30. Epub 2013/11/15. doi: 10.1093/bioinformatics/btt656. PubMed PMID: 24227677.

46. Feng J, Meyer CA, Wang Q, Liu JS, Shirley Liu X, Zhang Y. GFOLD: a generalized fold change for ranking differentially expressed genes from RNA-seq data. Bioinformatics. 2012;28(21):2782–8. Epub 2012/08/28. doi: 10.1093/bioinformatics/bts515. PubMed PMID: 22923299.

47. Love MI, Huber W, Anders S. Moderated estimation of fold change and dispersion for RNA-seq data with DESeq2. Genome Biol. 2014;15(12):550. Epub 2014/12/18. doi: 10.1186/s13059-014-0550-8. PubMed PMID: 25516281; PubMed Central PMCID: PMCPMC4302049.

48. Chen S, Zhou Y, Chen Y, Gu J. fastp: an ultra-fast all-in-one FASTQ preprocessor. Bioinformatics. 2018;34(17):i884–i90. Epub 2018/11/14. doi: 10.1093/bioinformatics/bty560. PubMed PMID: 30423086; PubMed Central PMCID: PMCPMC6129281.

49. Langmead B, Salzberg SL. Fast gapped-read alignment with Bowtie 2. Nat Methods. 2012;9(4):357–9. Epub 2012/03/06. doi: 10.1038/nmeth.1923. PubMed PMID: 22388286; PubMed Central PMCID: PMCPMC3322381.

50. Tarasov A, Vilella AJ, Cuppen E, Nijman IJ, Prins P. Sambamba: fast processing of NGS alignment formats. Bioinformatics. 2015;31(12):2032–4. Epub 2015/02/24. doi: 10.1093/bioinformatics/btv098. PubMed PMID: 25697820; PubMed Central PMCID: PMCPMC4765878.

51. Zhang Y, Liu T, Meyer CA, Eeckhoute J, Johnson DS, Bernstein BE, et al. Model-based analysis of ChIP-Seq (MACS). Genome Biol. 2008;9(9):R137. Epub 2008/09/19. doi: 10.1186/gb-2008-9-9-r137. PubMed PMID: 18798982; PubMed Central PMCID: PMCPMC2592715.

52. Heinz S, Benner C, Spann N, Bertolino E, Lin YC, Laslo P, et al. Simple combinations of lineage-determining transcription factors prime cis-regulatory elements required for macrophage and B cell identities. Mol Cell. 2010;38(4):576–89. Epub 2010/06/02. doi: 10.1016/j.molcel.2010.05.004. PubMed PMID: 20513432; PubMed Central PMCID: PMCPMC2898526.

53. Ramírez F, Dündar F, Diehl S, Grüning BA, Manke T. deepTools: a flexible platform for exploring deep-sequencing data. Nucleic Acids Res. 2014;42(Web Server issue):W187-91. Epub 2014/05/07. doi: 10.1093/nar/gku365. PubMed PMID: 24799436; PubMed Central PMCID: PMCPMC4086134.

54. Martin M. Cutadapt removes adapter sequences from high-throughput sequencing reads. EMBnetjournal; Vol 17, No 1: Next Generation Sequencing Data AnalysisDO - 1014806/ej171200. 2011.

55. Yu G, Wang LG, He QY. ChIPseeker: an R/Bioconductor package for ChIP peak annotation, comparison and visualization. Bioinformatics. 2015;31(14):2382–3. Epub 2015/03/15. doi: 10.1093/bioinformatics/btv145. PubMed PMID: 25765347.

56. Quinlan AR, Hall IM. BEDTools: a flexible suite of utilities for comparing genomic features. Bioinformatics. 2010;26(6):841–2. Epub 2010/01/30. doi: 10.1093/bioinformatics/btq033. PubMed PMID: 20110278; PubMed Central PMCID: PMCPMC2832824.

57. Wang M, Kong L. pblat: a multithread blat algorithm speeding up aligning sequences to genomes. BMC bioinformatics. 2019;20(1):28. Epub 2019/01/17. doi: 10.1186/s12859-019-2597-8. PubMed PMID: 30646844; PubMed Central PMCID: PMCPMC6334396.

